# Identification of potential treatments for COVID-19 through artificial intelligence-enabled phenomic analysis of human cells infected with SARS-CoV-2

**DOI:** 10.1101/2020.04.21.054387

**Authors:** Katie Heiser, Peter F. McLean, Chadwick T. Davis, Ben Fogelson, Hannah B. Gordon, Pamela Jacobson, Brett Hurst, Ben Miller, Ronald W. Alfa, Berton A. Earnshaw, Mason L. Victors, Yolanda T. Chong, Imran S. Haque, Adeline S. Low, Christopher C. Gibson

## Abstract

To identify potential therapeutic stop-gaps for SARS-CoV-2, we evaluated a library of 1,670 approved and reference compounds in an unbiased, cellular image-based screen for their ability to suppress the broad impacts of the SARS-CoV-2 virus on phenomic profiles of human renal cortical epithelial cells using deep learning. In our assay, remdesivir is the only antiviral tested with strong efficacy, neither chloroquine nor hydroxychloroquine have any beneficial effect in this human cell model, and a small number of compounds not currently being pursued clinically for SARS-CoV-2 have efficacy. We observed weak but beneficial class effects of β-blockers, mTOR/PI3K inhibitors and Vitamin D analogues and a mild amplification of the viral phenotype with β-agonists.

---

SARS-CoV-2, a novel coronavirus, emerged in December 2019 in Wuhan, China. Over the following months the resulting disease, subsequently named COVID-19, spread across the rest of the world and was declared a pandemic by the World Health Organization on March 11th, 2020. As of submission, there are over 2.2M reported infections worldwide and over 150,000 reported deaths, with an anticipated world wide death toll in the millions and an economic impact sufficient to cause a worldwide recession or depression^1,2^. There are currently no vaccines available and the only available therapies are supportive or based largely on anecdotal work.

To rapidly identify and prioritize potential treatments of COVID-19, we adapted our deep learning-enabled drug discovery platform to evaluate a library of FDA-approved drugs, EMA-approved drugs or compounds in late stage clinical trials for modulation of the effect of SARS-CoV-2 on human cells. Through unbiased image analysis of cytological structures of perturbed cells (phenomics), profiles can be rapidly created for use as the basis of chemical suppressor screens^3,4^. Such profiles can be generated to model many diseases across multiple therapeutic areas, including: inflammatory disease (cytokines), genetic diseases (siRNAs or CRISPR), or infectious disease with the live unmodified SARS-CoV-2 virus. This method can be deployed rapidly for high throughput phenotypic screening in any relevant cell types - including human - without development of disease-specific tools such as antibodies, reporters, or surrogate cell lines. This technique can detect traditional inhibitors of viral replication, but also novel or unexpected signals that could come about through disrupting the interactions of the virus with the host or modulating cytokine signalling.

To identify a human SARS-CoV-2 model suitable for high throughput screening, we infected monolayers of normal human renal cortical epithelial cells (HRCE), as well as normal human bronchial epithelial cells and Caco-2 cells, then fixed, stained, and imaged cells at 96 hours post infection as described previously (**Figure 1a** and **Supplemental Figure 1a,b**)^4^. We also infected African green monkey kidney epithelial cells (Vero) as a control condition (**Supplemental Figure 1c**). To evaluate the specificity of the phenotypic profile to the impact of the virus we compared the effects of active SARS-CoV-2 to inactive ultraviolet-irradiated (UV-IR) and mock preparations of SARS-CoV-2 on cells. Images were processed using a proprietary deep learning neural network to generate high-dimensional featurizations of each image for the identification of distinct phenotypic profiles. Both HRCE and Vero cells demonstrated robust phenotypes compared to both mock and irradiated controls (Z-factors: HRCE active virus vs mock, 0.432; HRCE active virus vs irradiated, 0.438; Vero active virus vs mock, 0.803; Vero active virus vs irradiated, 0.827) (**Figure 1b**). Notably, the phenomics assay achieved the aforementioned, highly separable assay window in HRCEs without significant changes in cell count at this time point compared to both mock and UV-IR controls. HRCEs were selected for further screening due to their disease relevance and robust disease-specific phenotype^5^.

**Figure 1:**
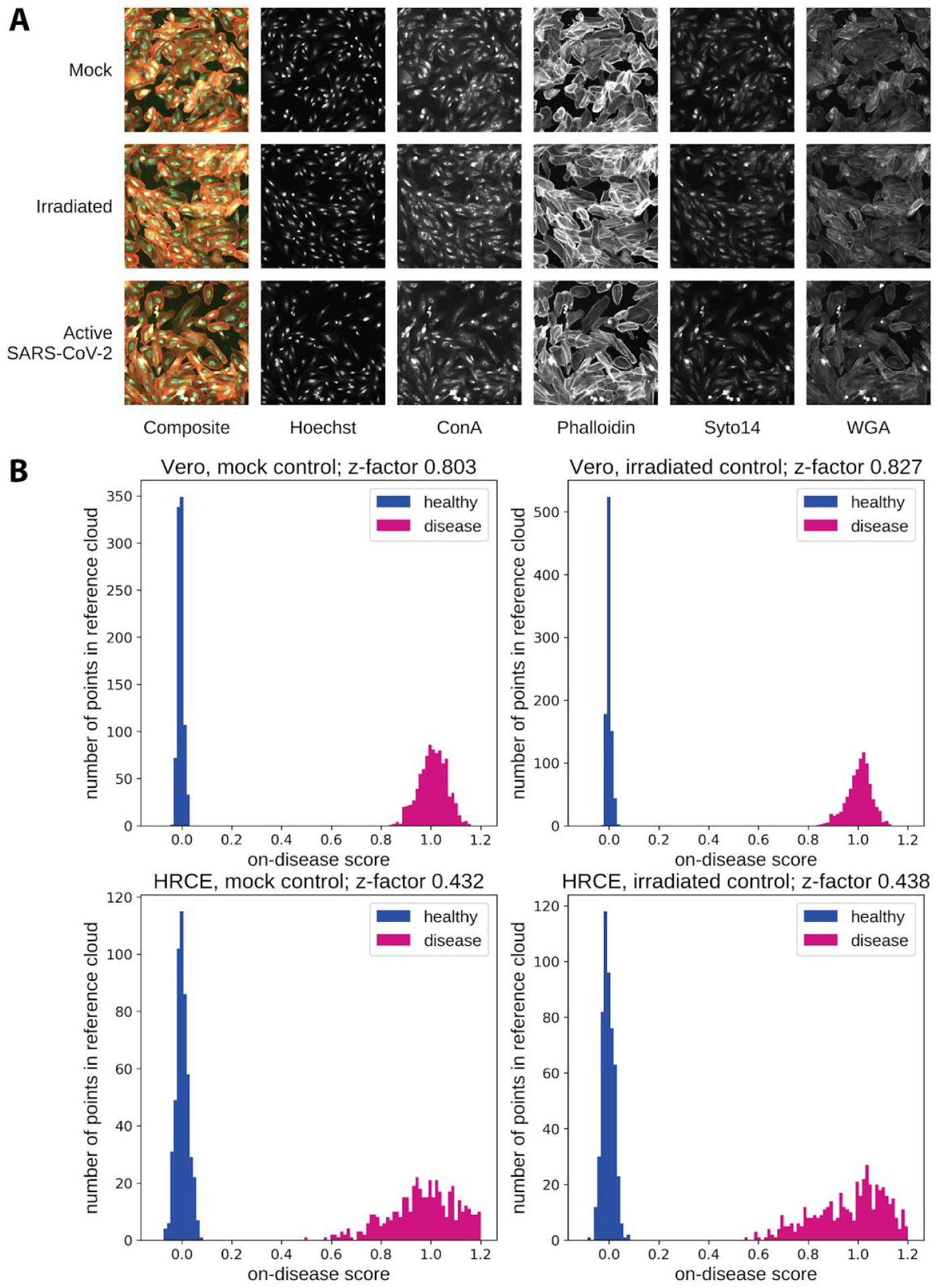
Analysis of SARS-CoV-2 infected cells identifies a screenable profile through deep learning. **a**) Representative images of HRCE from SARS-CoV-2 infected and control wells, including both a composite image and individual channels for each unique stain. **b**) Distribution histograms of a test split of the control groups. The X-axis represents the on-disease score and the Y-axis is the number of wells. SARS-2-CoV treated wells (pink) and control wells (blue) by control type (left column mock, right column UV irradiated virus) and cell type (top row Vero, bottom row HRCE).

Chemical Suppressor screens were conducted by treating HRCE cells in six half-log doses with six replicates per dose for each compound approximately two hours after cell seeding. At 24 hours post seeding, cells were infected with SARS-CoV-2 at a MOI of 0.4 and incubated for 96 hours until fixation, staining and imaging. Following imaging, compounds were evaluated for their impact on the morphological profile of infection as defined by the difference between the active virus and the mock condition (‘on-disease score’) plus their overall impact on other elements of cell morphology not associated with the disease model (‘off-disease score’) (**Figure 2a, Supplemental Figure 2**). In vero cells, chloroquine, lopinavir, and remdesivir exhibited disease score suppression activity at IC50 values similar to what has been previously established for SARS-CoV-2 in cytopathic effect (CPE) and viral N-protein expression assays^6,7^. A proprietary hit score algorithm based on the comparative on-disease vs off-disease score in dose-response was used to determine efficacy of compounds (**Figure 2b)**.

**Figure 2:**
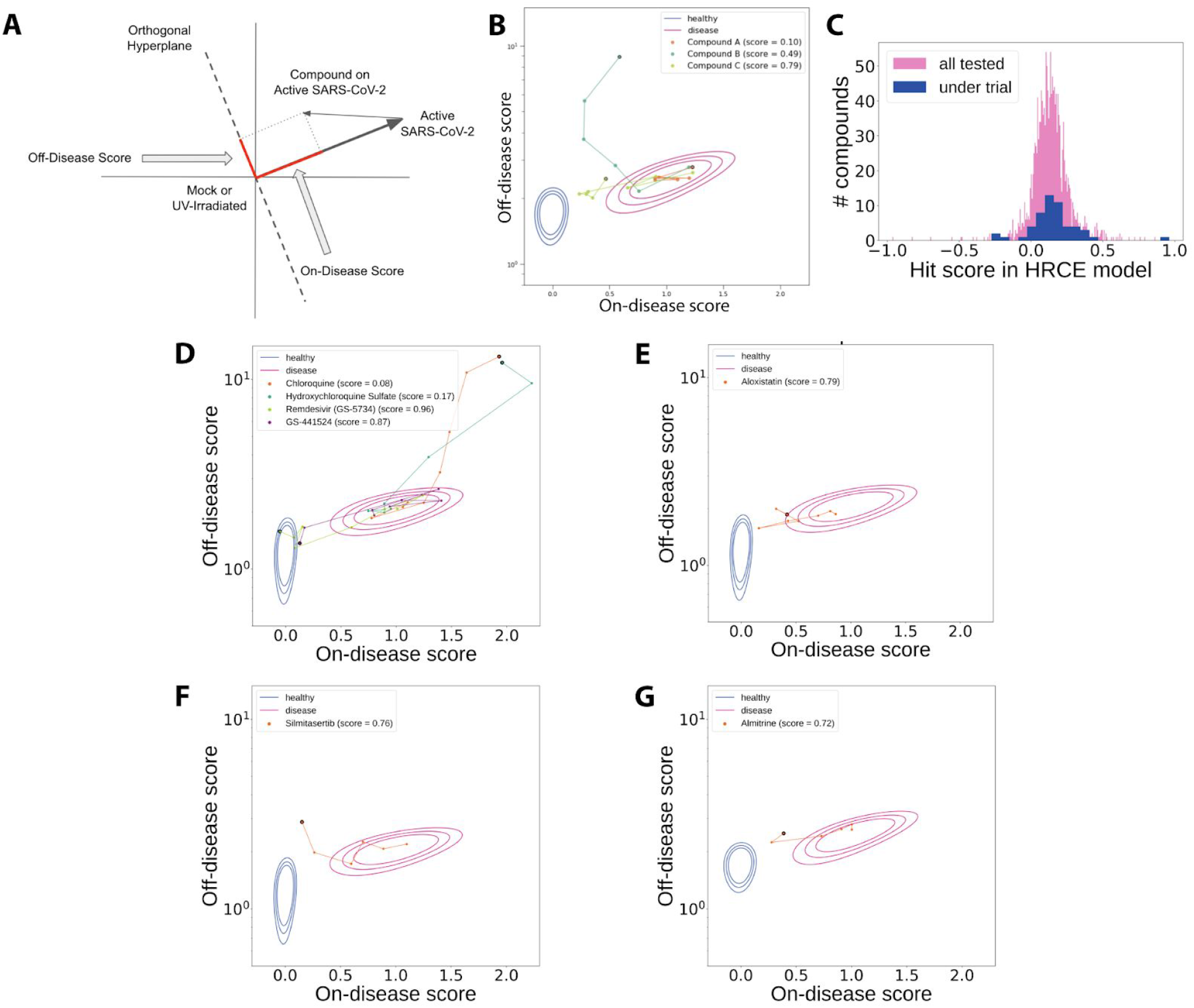
High-dimensional activity of molecules in HRCE infected with SARS-CoV-2. **a**) High-dimensional image embeddings were deconvolved into measures of on-disease and off-disease morphology by projecting onto both the vector between the mock condition and the active SARS-CoV-2 condition as well as the orthogonal hyperplane to this vector. **b**) Mock control populations (‘healthy’) and SARS-CoV-2-infected populations (‘disease’) are shown. Expanding contours denote the 50th, 90th, and 99th percentiles of the populations. Glyphs represent the median of six technical replicates. Compounds were scored using a proprietary algorithm that prioritizes low on-disease scores and penalizes high off-disease scores. Example compounds with increasing hit scores and prototypical trajectories are shown. Connected line segments represent the trajectory between increasing doses for each compound, culminating in the large glyph which represents the highest dose tested for each. **c**) Distribution of hit scores for all molecules tested (pink) and molecules currently in clinical investigation (blue). **d**) Remdesivir and its parent nucleoside demonstrated strong efficacy in this human model, whereas chloroquine and hydroxychloroquine showed no benefit. All three of **e**) Aloxistatin, **f**) Silmitasertib, and **g**) Almitrine demonstrated strong efficacy without inducing off-disease morphological changes.

The vast majority of compounds currently reported as under evaluation in human clinical trials for COVID-19 showed no or weak on-disease efficacy (**Figure 2c**)^8^. However, remdesivir (as well as its parent nucleoside GS-441524) demonstrated strong repeatable efficacy in our model (**Figure 2d** and **Supplemental Figure 3**). Of particular interest, chloroquine and hydroxychloroquine showed no benefit in the human cell model; however, they showed weak benefit in Vero cells, albeit with very high off-disease activity (**Figure 2d** and **Supplemental Figure 4**).

Next we report on three compounds not currently in trials for COVID-19 which demonstrated moderate to strong effects in our model. The two clinical-stage assets were added to our library based on *in silico* or *in vitro* analyses suggesting their potential utility.

Aloxistatin (E64d) is an irreversible cysteine protease inhibitor initially developed for muscular dystrophy and advanced through Phase 3 in Japan^9^ (**Figure 2e**). Recent studies have confirmed that the cysteine protease cathepsin L is required for SARS-CoV-2 viral entry, and aloxistatin treatment reduced cellular entry of SARS-CoV-2 pseudovirions by 92.3%^10^.

Silmitasertib (CX-4945) is a selective inhibitor of protein kinase casein kinase 2 (CK2) currently in clinical development in oncology^11^ (**Figure 2f**). CK2 was recently suggested as a therapeutic target for SARS-CoV-2 based on large scale interactome data and hypothesized to promote an antiviral cellular state by reducing turnover of stress granules^12^. Alternatively, results from SARS-CoV-1 suggest that CK2 inhibition may prevent nucleocapsid shuttling from the nucleus to the cytoplasm^13^.

Almitrine is a respiratory stimulant used in acute respiratory failure such as chronic obstructive pulmonary disease (COPD) that is available in Europe, but not currently available in the US^14^ (**Figure 2g**). Recent structure-based computational studies hypothesized that almitrine binds to the SARS-CoV-2 main viral proteinase (Mpro, or 3CLpro), an important viral target given the necessity of this protein for viral replication^15–17^. Our results demonstrate for the first time activity of almitrine in human cells infected with SARS-CoV-2, suggesting almitrine may represent a novel SARS-CoV-2 protease-targeting antiviral.

Further, we have identified chemical classes of compounds which, across the class, display weak but consistent rescue, and are reasonably well-tolerated in humans, offering the potential to be elements of combination therapies.

The most significantly enriched class of molecules identified were beta-adrenergic receptor antagonists with 9 of 32 molecules demonstrating above threshold activity (**Supplemental Figure 5a**, *P* < 0.0001). Among the most active molecules in the class were carvedilol and celiprolol; both are molecules with combined beta and alpha adrenergic blockade, properties that are thought to enhance the effectiveness of these drugs in patients with combined heart failure and COPD^18^. An association between beta adrenergic blocker use and SARS-COV-2 has not been previously demonstrated, but given the elevated risk of mortality among patients with preexisting heart disease^19^, it is intriguing to consider whether subclasses may offer enhanced benefit for these patients.

A second mechanistic class that demonstrated enrichment was PI3K/mTOR. Sirolimus and its derivative compounds, and two of three PI3K inhibitors demonstrated above threshold activity (**Supplemental Figure 5b**, *P* < 0.0001). Therapeutic benefit of mTOR modulating agents in SARS-CoV-2 was hypothesized based on human protein-protein interactome results suggesting enriched interactions of viral proteins with human targets of mTORC1^12^. Similarly, a systems computational pharmacology approach predicted sirolimus as a putative therapeutic based on SARS-CoV-2 protein network mapping^20^. Notably, inhibition of mTOR with sirolimus reduced MERS-CoV infection by ~60% in cell-based studies^21^. Furthermore, diabetes represents a second major risk factor for severe SARS-CoV-2, and given dysregulation of mTOR signaling in the diabetic state, it is appealing to consider these results may reveal a mechanistic basis for adverse outcomes in these patients^19^.

A third mechanistic class that demonstrated significant enrichment among actives in the screen was vitamin D receptor agonists (**Supplemental Figure 5c**, *P* < 0.0001). While the role of vitamin D in immunity is well established^22,23^ and vitamin D supplementation is subject of an ongoing investigational study^24^, limited evidence supports a beneficial role for vitamin D in prevention or resolution of SARS-CoV-2 infection. A large meta-analysis of acute respiratory tract infection found that supplementation was beneficial overall, but most protective in patients with baseline vitamin D deficiency^25^.

Lastly, several notable mechanistic classes demonstrated lack of any activity or were significantly enriched among molecules that trended toward amplifying the disease phenotype. ACE inhibitors and angiotensin receptor blockers (ARBs) have been the subject of interest given their widespread use and reported viral entry into the cell via the ACE2 receptor^26^. These classes showed no significant activity. However, beta-adrenergic receptor agonists such as albuterol and formoterol commonly administered as bronchodilators led to a statistically significant amplification of the SARS-CoV-2 phenotypes (**Supplemental Figure 5d**, *P* <0.0001) and may warrant caution in patients with COVID-19.

Together, these data show for the first time a high-dimensional analysis of more than 1,660 approved drugs in a human cellular model of SARS-CoV-2 infection. Although this in vitro screen represents data from only a single human cell type, HRCE, this method is likely broadly applicable to other primary human cell models. These data may be useful as the basis for prioritizing further work given constrained resources and serve to generate hypotheses for other researchers in this rapidly progressing field. In our case, this data will serve as the basis for ongoing efforts to identify a cocktail of additive or synergistic, orally available, rapidly manufacturable, and safe FDA-approved drugs with robust activity for further study in animals or humans as a stop-gap to a vaccine, and more broadly utilizable than remdesivir. Finally, from signing an agreement with our BSL3 partner facility to submitting these data required only 4 weeks. With some optimization, this type of work could potentially be conducted in under 3 weeks and across much larger chemical libraries, suggesting that this broad approach—which is able to identify in vitro signals of efficacy across a wide variety of mechanisms—could be a robust tool for rapid therapeutic discovery in future pandemics.

## Supplemental Figures

**Supplemental Figure 1:**
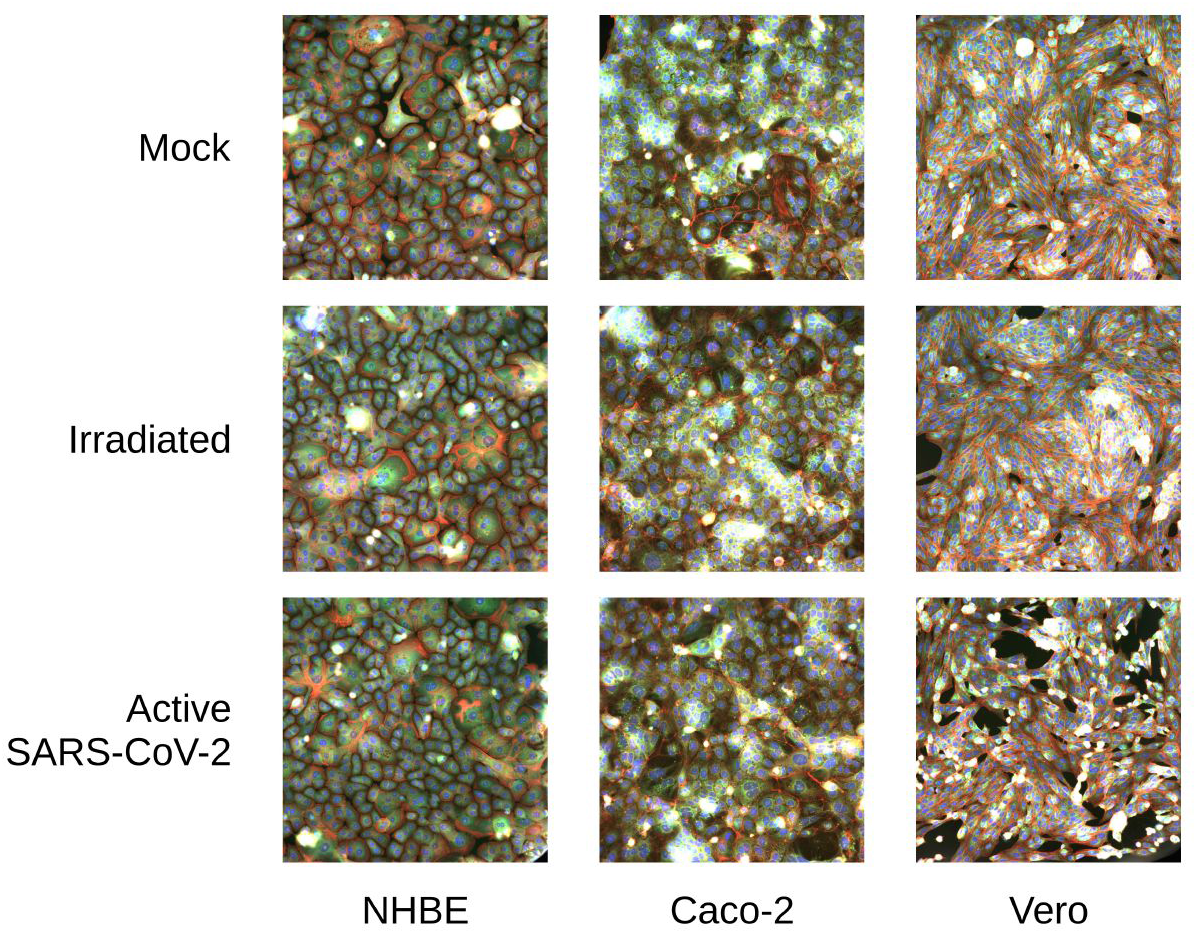
Analysis of additional cell types treated with SARS-CoV-2. Representative images of **a**) NHBE, **b**) Caco2 and **c**) Vero cells in each control and active SARS-CoV-2 treated representative wells.

**Supplemental Figure 2:**
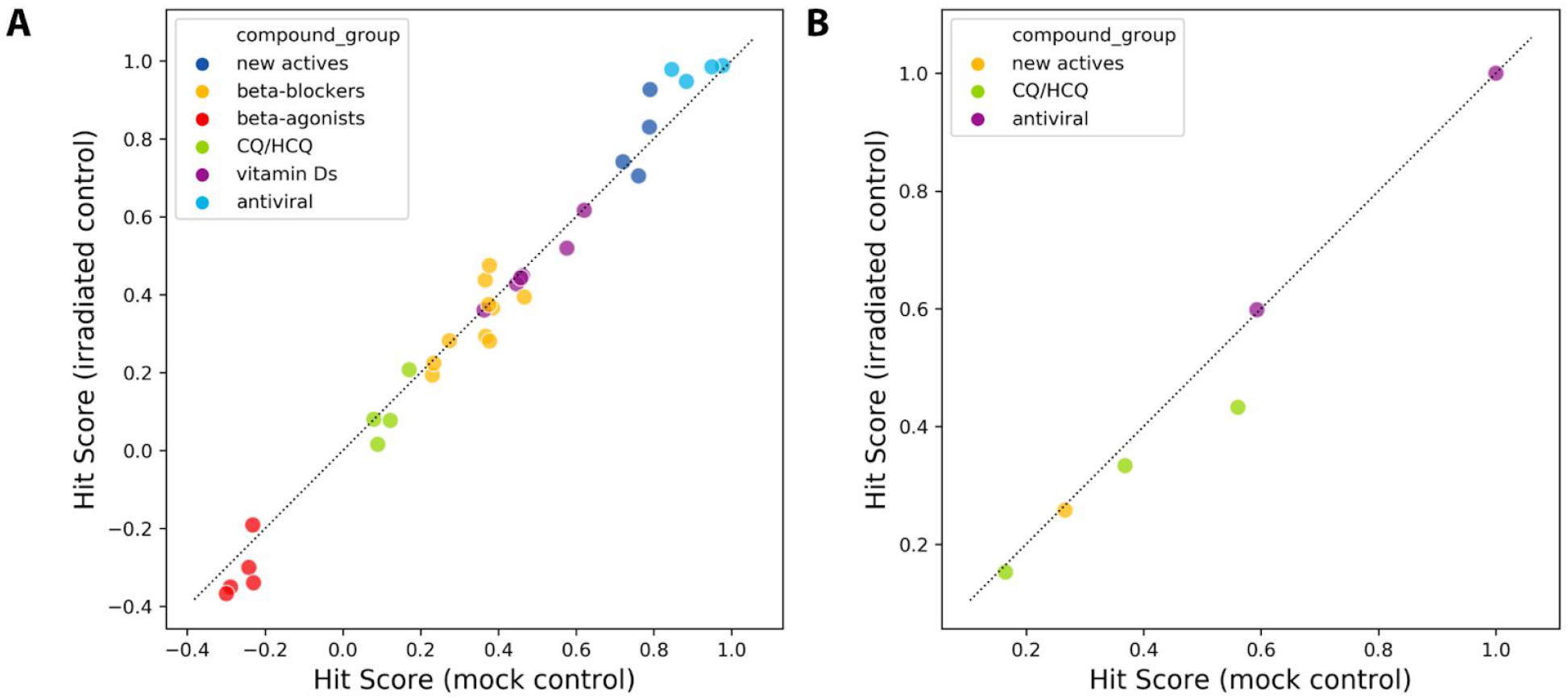
Evaluation of select hit scores against mock and irradiated control groups in HRCE and VERO cells infected with SARS-CoV-2. In addition to evaluation against the mock control group, all compounds were also evaluated against the UV irradiated control condition. The resulting hit scores demonstrate high correlation independent of control condition. Shown here are many of the compounds highlighted in the text in both **a)** HRCE and **b)** Vero cells. Compounds groups shown here represent: (antiviral) Remdesivir (GS-5734), GS-441524; (new actives) Aloxistatin, CX-4945, Almitrine; (CQ/HCQ) Chloroquine, Thymoquinone, Hydroxychloroquine Sulfate; (beta-blockers) Bupranolol, Sotalol, Arotinolol, Alprenolol, Pindolol, Bevantolol, Propranolol, Betaxolol, Bopindolol, Atenolol; (vitamin Ds) Calcipotriene, Paricalcitol, Doxercalciferol, Calcifediol, Tacalcitol, Secalciferol; (beta-agonists) Formoterol, Levalbuterol, Zilpaterol, Indacaterol, Albuterol.

**Supplemental Figure 3:**
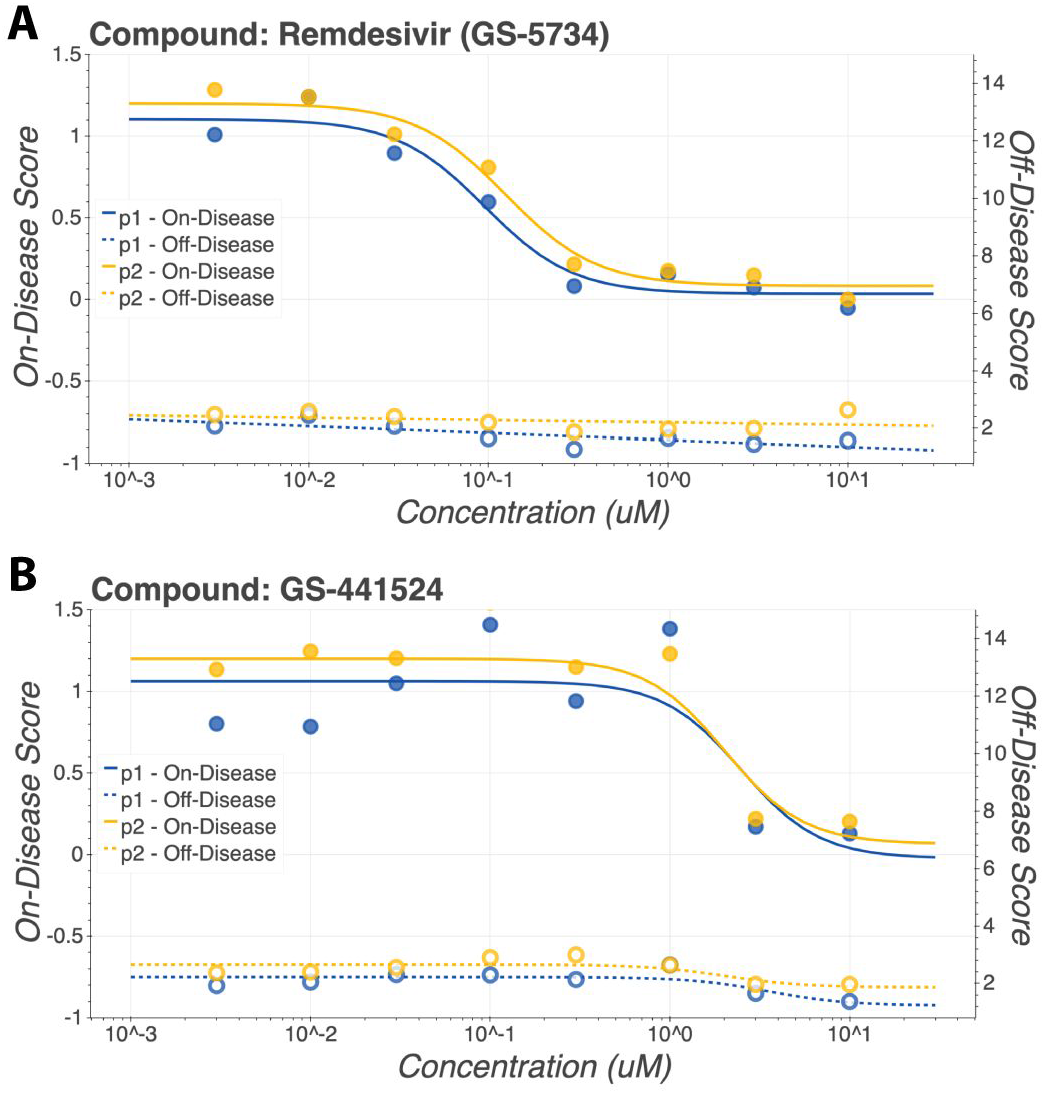
Compound response curves of Remdesivir and GS-441524 in HRCE infected with SARS-CoV-2. Compound response curves were computed for both on-disease scores and off-disease scores. On-disease score curves are shown with solid lines and glyphs while off-disease score curves are shown with dashed lines and unfilled glyphs. **a**) Remdesivir and its parent nucleoside **b**) GS-441524 show consistent behavior across experimental replicates.

**Supplemental Figure 4:**
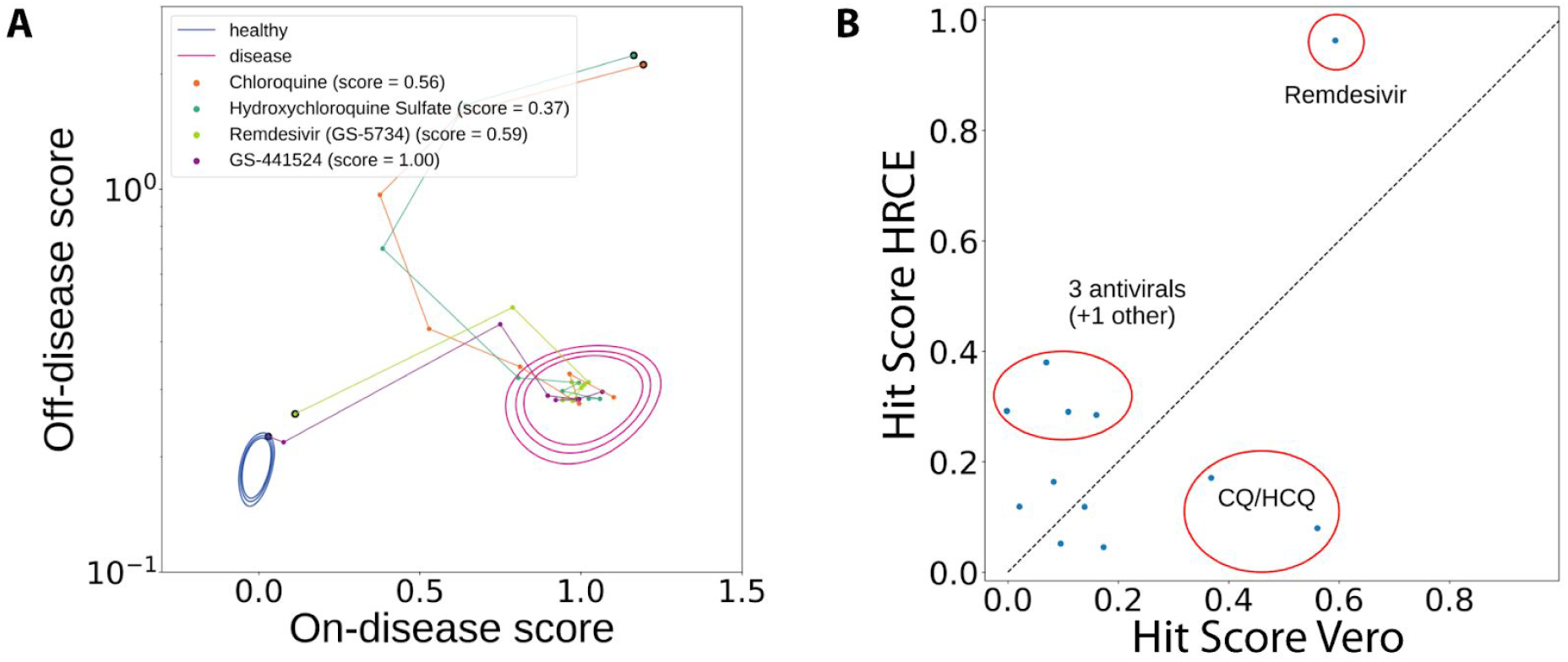
Profiles of various compounds vary across HRCE and Vero cells. **a**) Chloroquine, Hydroxychloroquine, Remdesivir and its parent nucleoside dose response in Vero cells. **b**) A comparison of various hit scores between experiments in HRCE and Vero cells.

**Supplemental Figure 5:**
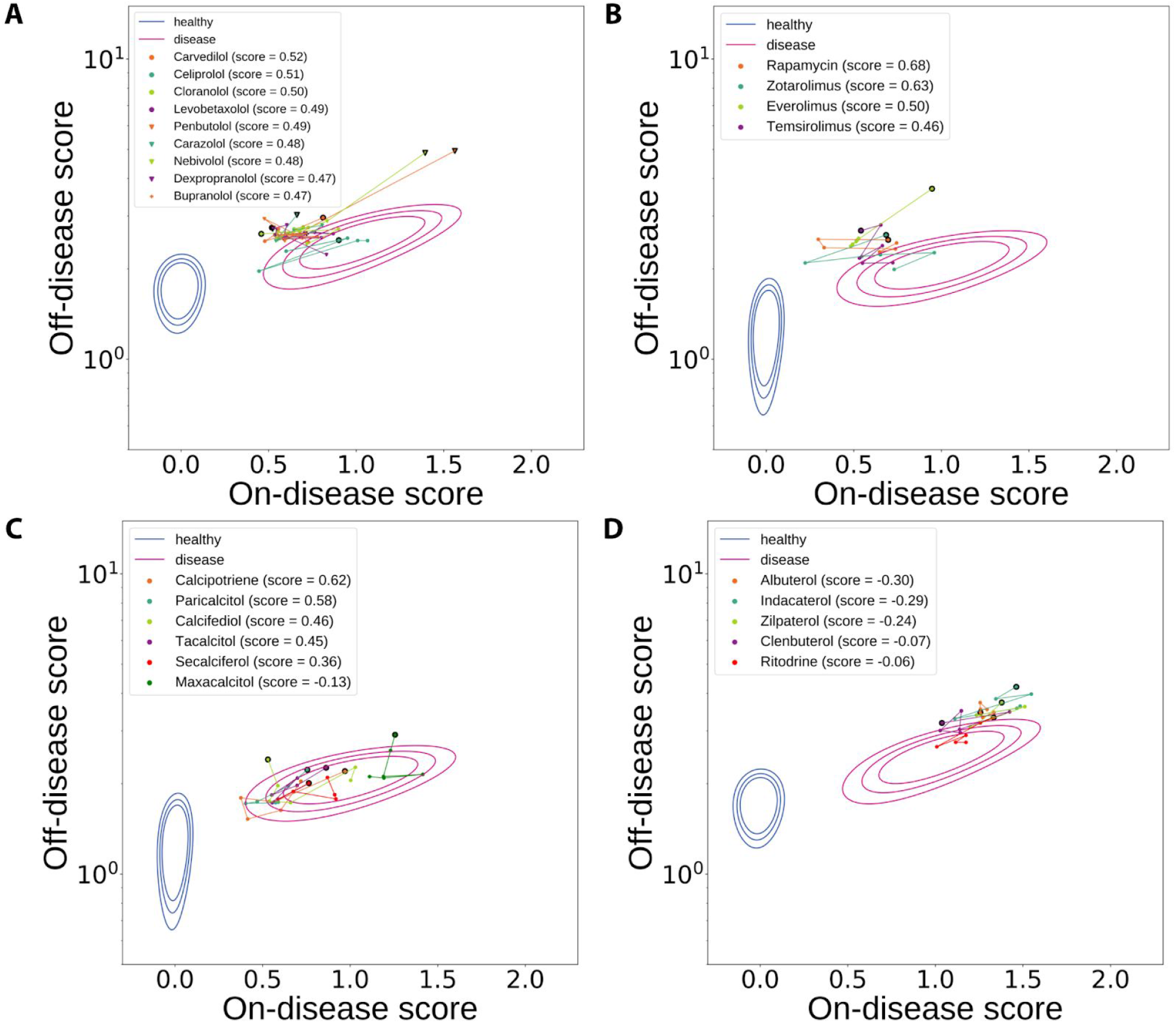
Significant class activity of molecules in HRCE infected with SARS-CoV-2. **a**) 9 of 32 Beta-adrenergic receptor antagonists show above-threshold activity. **b**) Sirolimus and its derivative compounds show above-threshold activity. **c**) Multiple vitamin D receptor agonists show above-threshold activity. **d**) beta-adrenergic receptor agonists led to a statistically significant amplification of the SARS-CoV-2 phenotypes.

## Acknowledgements

Charles Baker, Celeste Belletire, Kathleen Bennett, Dan Berger, Katie Brown, Nicholas Campbell, Joe Carpenter, Hannah Cook, Ryden Crowther, Ashley Dagley, Eric Fish, Briana Freshner, Jann Gardner, Mike Genin, Alex Gosch, Matthew Guthrie, Sharath Hegde, Kie Hoon-Jung, Jamie Jarvis, Dylan Knutson, Ben Mabey, Travis Martin, Catherine Maxwell, Brandon Mendivil, John Morrey, Chad Nielsen, Scott Nielsen, Kelsey Payne, Alexis Ramos, Katy Van Pelt, Parker Weber, Nate Wilkinson, AJ Williams, Marika Xydes and all employees of Recursion and Utah State University who have been working under extraordinary circumstances, home-schooling and socially-distancing, yet making it happen all the way through.

## Supplemental Methods

### Data availability

Nearly 300,000 5-channel immunofluorescence images underlying this report will be made publicly available at RxRx.ai along with deep learning embeddings of all images under the title “RxRx19a Dataset.”

### SARS-CoV-2 propagation and controls

The USA-WA1/2020 strain of SARS-CoV-2 was propagated in Vero 76 cells (ATCC CRL-1587). Cells were grown in standard tissue culture flasks (60% confluence) and were infected at a multiplicity of infection (MOI) of 0.001, in Eagle’s Minimum Essential Medium (EMEM) + 2% Fetal Bovine Serum (FBS) and 50 g/mL gentamicin, incubated at 37°C with 5% CO2 for 5 days. Supernatants containing virus were removed from these cultures, spun down to remove cellular debris and stored at −80°C until use. Viral titers were determined through standard tissue culture infectious dose 50% (TCID50) methods, where cytopathic effect (CPE) on Vero 76 cells was measured by visual observation under a light microscope.

To create a suitable control with inactivated virus, SARS-CoV-2 was irradiated with a UV lamp for 20 minutes. Viral inactivation in this sample was verified using visual CPE on Vero cells, where undetectable level of active virus was observed. An additional “mock” control was created using conditioned media preparations generated from uninfected Vero 76 cells grown in 2% FBS in EMEM for five days. Cellular debris were removed through centrifugation and the supernatants were frozen at −80°C until use.

All experiments using SARS-CoV-2 were performed at Utah State University using Biosafety Level 3 (BSL-3) containment procedures.

### Cell culturing, infection and compound addition

Vero cells were acquired from ATCC (CCL-81) and propagated at 37°C with 5% CO2 in EMEM and supplemented with 10% FBS. Normal Human Renal Cortical Epithelial cells (HRCE) were acquired from Lonza (CC-2554) and propagated at 37°C with 5% CO2 in EpiCM, (ScienCell # 4101) supplemented with Epithelial Cell Growth Supplement (EpiCGS, ScienCell #4152).

Caco-2 cells (HTB-37) were acquired from ATCC and propagated at 37°C with 5% CO2 in EMEM + 10% FBS. Normal Human Bronchial Epithelial cells (NHBE) were acquired from Lonza (CC-2540) and propagated according to manufacturer’s recommendations. Cells were harvested from active culture and were seeded at 750 cells per well in 1536-well microplates (Greiner Bio One, 789866) using a Multidrop Combi (Thermo Fisher). Cells were allowed to settle and attach for two hours, prior to compound addition using an Echo 555 Acoustic Liquid Handler (Labcyte). Most non-reference compounds were added in six half-log doses from 0.01 to 3.0 uM. Most reference compounds were added in eight half-log doses from 0.003 to 10.0 uM. Hydroxychloroquine Sulfate, Chloroquine, and Thymiquinone were dispensed in eight half-log doses from 0.03 to 100.0 uM; Bafilomycin A1 was dispensed in six half-log doses from 0.001 to 0.3 uM. Plates were then transported to the BSL3 facility and virus was added to cells approximately 20 hours after compound addition and incubated for the duration of the experiment (48 hrs or 96 hrs) at 37°C with 5% CO2. Vero cells were infected at an MOI of approximately 0.08 in EMEM supplemented with 2% FBS. HRCEs, CaCo2s and NHBEs were infected at an MOI of approximately 0.4 in EMEM supplemented with 2% FBS.

### Staining and imaging

Samples in 1536-well microplates were stained using a modified cell painting protocol^4^. Briefly, cells were fixed in 5% paraformaldehyde, permeabilized with 0.25% Triton X100, and stained with Hoechst 33342 (Thermo), Alexa Fluor 568 Phalloidin (Thermo), Alexa Fluor 555 Wheat germ agglutinin (Thermo), Alexa Fluor 488 Concanavalin A (Thermo), and SYTO 14 (Thermo) for 35 minutes at room temperature and then washed and stored in HBSS+0.02% sodium azide as previously described^4^. All reagent additions and washes were conducted using a Centrifugal Blue^®^ Washer (BlueCat Bio).

Plates were imaged using Image Express Micro Confocal High-Content Imaging System (Molecular Devices) microscopes in widefield mode with 20X objectives. Four sites per well were acquired with 5 channels per site. The following bandpass filters were used to visualize the five channels: FF409/493/573/652, FF459/526/596, FF01-432/515/595/730-25, FF01-475/543/702, and FF01-600/37/25.

### Feature extraction

Five-channel images were embedded into a 1024-dimensional feature space using a proprietary deep convolutional neural network trained on a 125,000+ image cellular imaging dataset Recursion has previously made public at RxRx.ai^27^. Post-processing of the extracted features included normalization to remove inter-plate variance, PCA to reduce the feature space, and anomaly detection to remove outliers from the active SARS-CoV-2, UV inactivated, and mock conditions. Featurization efficiency for representing infection was verified with a logistic regression classifier.

The vector pointing between the barycenters of the mock and active SARS-CoV-2 virus treated conditions was computed, and the extracted features were decomposed into the signed scalar projection (the on-disease score) and the scalar rejection (the off-disease score) with respect to this vector. These scores were normalized so that the mean disease score was 0 for the mock condition and 1 for the active SARS-CoV-2 virus condition (**Supplemental Table 1**). Separation of the mock and active SARS-CoV-2 virus conditions along the disease axis was verified by Z-factor.

### Hit Selection

Each compound dose was assigned a hit score using a proprietary algorithm that prioritizes low on-disease scores and penalizes high off-disease scores, with 1 being the highest possible hit score. For each compound, the two best-scoring doses were used to compute an overall hit score (**Supplemental Table 2**). Hits were selected from individual compounds with a hit score above 0.6 and compound classes that consistently showed hit scores above 0.45 by a hypergeometric test with p-value less than 0.05.

### Code availability

The code underlying this report leverages proprietary algorithms for image processing, data standardization, outlier detection and compound efficacy scoring. As such the code underlying this report will not be made available. Instead, much of the output of these algorithms is provided in Supplemental Tables 1 and 2.

